# Environmental factors shape methionine metabolism in p16/MTAP deleted cells

**DOI:** 10.1101/313288

**Authors:** Sydney M. Sanderson, Peter Mikhael, Ziwei Dai, Jason W. Locasale

## Abstract

The co-deletion of a common tumor suppressor locus and neighboring metabolic gene is an attractive possible synthetic dependency of tumor suppression on metabolism. However, the general impact that these co-deletions have on metabolism, which also dependent on nutrient availability and the tissue of origin, is unknown. As a model to investigate this question, we considered a set of tissue-matched cancer cells with homozygous co-deletions in *CDKN2a* and *MTAP*, genes respectively encoding the most commonly deleted tumor suppressor p16 and an enzyme involved in methionine metabolism. A comparative metabolomics analysis revealed that while there exists a definite pan-cancer metabolic signature of MTAP-deletion, this signature was not preserved when cells were subjected to changes in the availability of methionine, serine, or cysteine, nutrients related to methionine metabolism. Notably, the heterogeneity exhibited by these cells in their responsiveness to nutrient availability dominated both MTAP status and tissue-of-origin. Furthermore, re-expression of MTAP exerted a modest effect on metabolism. Together these findings demonstrate that environmental factors, particularly nutrition and tissue identity, may overwhelm the genetic effects of collateral deletions of metabolic genes.

## Introduction

Metabolic phenotypes arise from a complex interaction between genes and environment. Determinants include the genomic encoding of metabolic genes and their sequence variants, transcriptional and allosteric regulation of metabolic enzyme activity, and nutrient availability, the latter of which is dependent on host nutrition and physiology as well as the local cellular microenvironment. This multitude of factors makes identifying contexts in which effectively targeting a specific aspect of metabolism challenging. Despite this complexity, the prospect of targeting metabolism for therapy is attractive due to the drugability of metabolic enzymes, and the numerous metabolic alterations observed between healthy and pathological conditions such as cancer. Nevertheless, principled strategies that define context-specific metabolic differences are desired.

One example of identifying these contexts considers the observation that genetic deletions of tumor suppressor genes are often accompanied by co-deletion of neighboring genes that encode metabolic enzymes. This approach exclusively relies on genetics to define metabolic pathway alterations that are specific to the genetic lesion of interest. For example, approximately 15% of cancers exhibit homozygous deletions of the *CDKN2a* locus, which encodes for the tumor suppressor p16, with 80-90% of these tumors also exhibiting concurrent deletion of a proximal gene, *MTAP*, that encodes for the methionine salvage enzyme methylthioadenosine phosphorylase (1, 2). A number of studies have investigated *MTAP* deletion as a possible collateral lethality and have identified vulnerabilities in this subset of cancers (3–7), with several investigating some of the effects of *MTAP* deletion on metabolism (8–10). However, a systematic comparison of the relative effects of *MTAP* deletion as it relates to other variables that shape metabolism is lacking.

The recycling of the essential amino acid methionine (i.e. methionine salvage) is an integral component of a metabolic network known as one-carbon metabolism (11). This network integrates nutrients such as glucose and dietary amino acids (including serine, cysteine, and methionine) for use in various biological processes such as phospholipid synthesis, redox maintenance, and production of substrates for methylation reactions (12–16). Changes to the availability of these nutrients through diet have been shown to exert major effects on the metabolism of this pathway; for example, dietary serine has been found to directly affect tumor growth (17, 18). Moreover, it has been demonstrated in many contexts that the same genetic alteration can have significantly different metabolic consequences depending on the tissue-of-origin (19, 20).

Utilizing a metabolomics approach (21–23), this current study quantifies the impact of *MTAP* deletion on metabolism in the context of cell type, tissue of origin, and the availability of nutrients related to methionine metabolism. We find that while *MTAP* deletion produces a defined pan-tissue metabolic signature, this signature is diminished upon consideration of the changes to metabolism that result from the availability of nutrients related to methionine and one-carbon metabolism. Furthermore, these changes vary widely across individual cell lines. Thus, upon considering other variables that shape metabolism, MTAP status alone appears to exert a relatively modest effect on cellular metabolism.

## Results

### MTAP status has a defined metabolic signature

MTAP uses the substrate methylthioadenosine (MTA) to allow for the recycling of methionine for the methionine cycle (Figure 1A). To begin to investigate its impact on metabolism, we first assembled a panel of 10 tissue-matched cancer cell lines, with each pair comprised of one cell line characterized by homozygous deletions of p16 and MTAP (MTAP ^-/-^) and the other exhibiting no alterations in this chromosomal locus (MTAP^+/+^); the presence or absence of MTAP protein was verified by immumoblotting (Figure 1B). Using liquid chromatography coupled with high resolution mass spectrometry (LC-HRMS), we analyzed the levels of over 200 metabolites between the cell lines in standard culture conditions to assess global metabolic profiles of each line (Figure S1). Extending on previous studies that found that MTAP status could predict differential MTA levels (8–10), we found that MTA was the most differentially abundant metabolite between the two groups, with MTAP-deleted lines exhibiting significantly higher levels of MTA compared to MTAP-expressing lines (Figure 1C).

**Figure 1:**
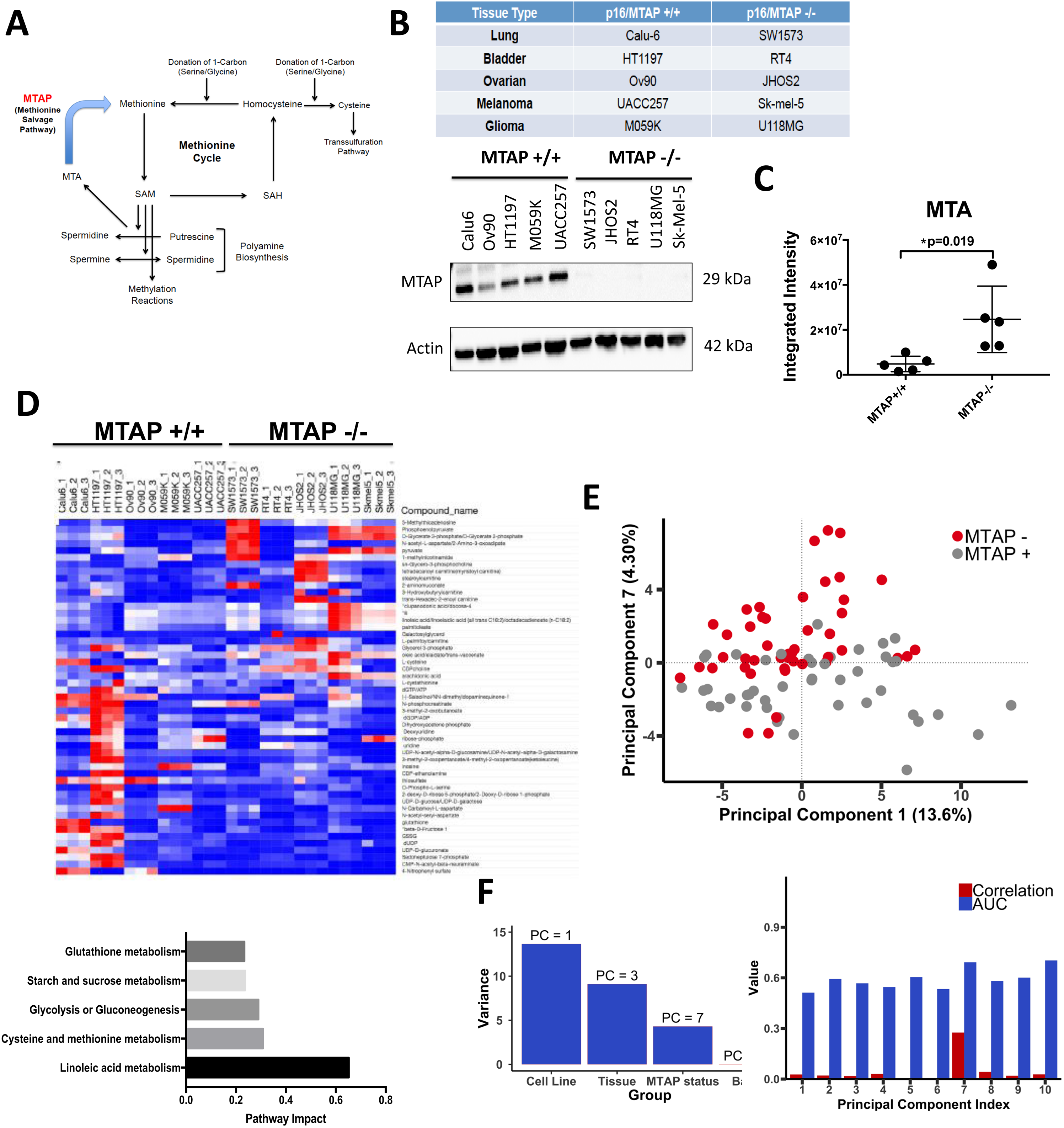
Cancer cell lines exhibiting homozygous deletion of MTAP show altered patterns of metabolite levels. (A) The methionine cycle. Methionine can be recycled from homocysteine via a donation from serine or glycine, or salvaged by MTAP via conversion of the polyamine biosynthesis byproduct MTA. (B) Cancer cell line panel of 10 lines from 5 different tissues, exhibiting either wild-type or homozygous deletion of p16/MTAP. Western blot validation of MTAP deletion. (C) Integrated intensity values (relative metabolite abundance) of MTA in MTAP+/+ vs MTAP-/- cell lines. P values were obtained from a student’s t-test.(D) Heat map of top 50 differential metabolites between MTAP+/+ and MTAP-/- cell lines. Top impacted pathways are indicated. (E) Principal component analysis of variance between MTAP+/+ and MTAP-/- groups. (F) Assignment of principal component that displays greatest variance between the indicated groups.

We next identified the top 50 differential metabolites between the MTAP^+/+^ and MTAP^-/-^ lines. When these metabolites were visualized using unsupervised hierarchical clustering, a pattern (i.e. MTAP “signature”) was evident (Figure 1D). Network-based pathway analysis (Methods) of these metabolites indicated that the metabolic profiles of MTAP^-/-^ lines differed from those of MTAP^+/+^ lines predominantly via differential activity of cysteine metabolism and glycolysis, consistent with our initial observations (Figure 1D).

We then performed principal component analysis (PCA) to determine the extent to which MTAP status segregated the metabolic profiles of the cell lines (Figure 1E). We found that the seventh principal component (PC7) best separated the two groups, accounting for 4.3% of the overall variance (Figure 1F). Comparatively, PC3 (accounting for ~8% of the variance) best separated the cell lines by tissue type. Of note, the majority of the variance was linked to the individual cell line (i.e. each cell line distinctly clustered from each other). These findings indicate that while *MTAP* deletion does induce a reproducible metabolic shift, heterogeneity between cell lines is a larger source of variation compared to tissue type and MTAP status (Figure 1F).

### Responsiveness to methionine availability is not predicted by MTAP status

In basal conditions, an MTAP-associated metabolic signature is readily apparent; however, metabolism has been shown to depend on nutrient availability and other environmental factors (12, 19, 24). Therefore, we next sought to determine whether MTAP-deleted lines would exhibit enhanced responsiveness to nutrient availability, particularly methionine restriction. We hypothesized that under these conditions, MTAP-associated metabolism would be affected due to an inability to salvage methionine from MTA. To test this, we cultured each cell line in either complete (100 uM methionine, or Met+) or methionine-restricted (3 uM methionine, or Met-) media for 24 hours, and used LC-HRMS to define the metabolic consequences of methionine restriction across the cell lines (Figure 2A, S2A).

**Figure 2:**
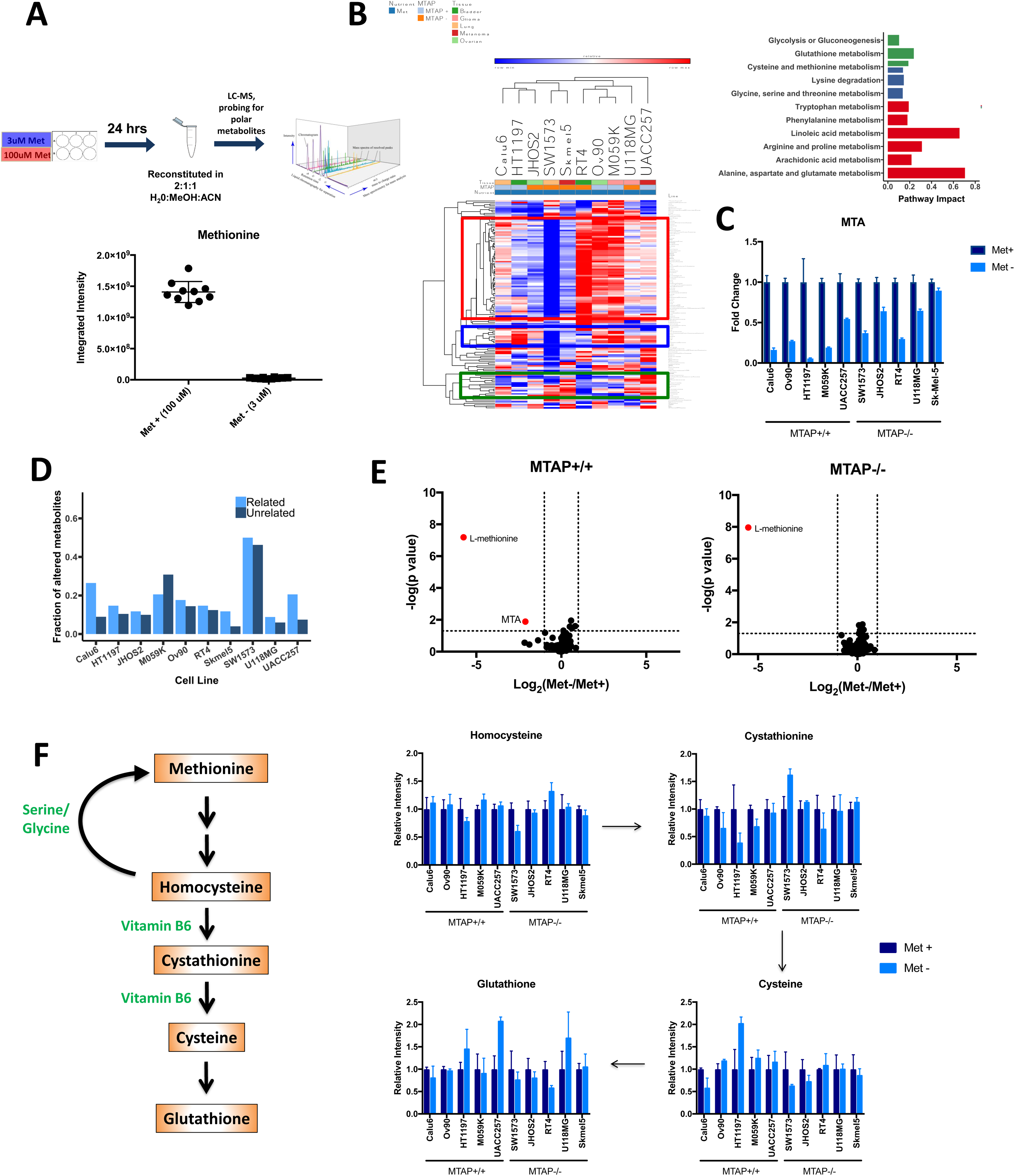
Responsiveness to methionine availability is heterogeneous and is not predicted by MTAP status. (A) Experimental set-up, and validation of methionine restriction. (B) Heat map of fold changes (FC) in global metabolite levels upon methionine restriction, hierarchically clustered by cell lines and metabolites. Top impacted pathways are indicated. (C) FC values of MTA metabolite levels either in complete (100uM Met, or Met+) or restricted (~3uM Met, or Met-) (D) Fraction of significantly altered (p<0.05) metabolites that are related (i.e. within 5 reactions) or unrelated to methionine. (E) Volcano plot of FC (Met-/Met+) values of metabolites, averaged across either MTAP+/+ or MTAP-/- groups. P values obtained using Student’s t-test. (F) Example of heterogeneous responsiveness to methionine restriction, as indicated by FC in metabolites involved in the transsulfuration pathway.

Unsupervised hierarchical clustering of both the individual samples and the fold changes of the metabolites revealed that the cell lines did not cluster by MTAP status or by tissue type (Figure 2B). Additionally, pathway analysis demonstrated that certain pathways tended to cluster together, such as fatty acid metabolism with tryptophan and phenylalanine metabolism, as well as pathways involved in one-carbon metabolism (including serine, glycine, cysteine, and methionine metabolism) (Figure 2B). Interestingly, MTA levels were significantly diminished in all except one (Sk-Mel-5) of the ten cell lines upon methionine restriction (Figure 2C). When we analyzed the number of significantly altered metabolites that were methionine-related (i.e. within 5 biochemical reactions) or methionine-unrelated (greater than 5 reactions away), MTAP^-/-^ lines were not found to exhibit a greater degree of altered methionine metabolism compared to MTAP^+/+^ lines upon methionine restriction (Figure 2D). Furthermore, PCA conducted for fold change values of each metabolite demonstrated that cell lines tended to cluster by MTAP status to a greater extent than by tissue of origin, but the majority of the variance was accounted for by cell line-to-cell line variability (Figure S2B), and cell lines that exhibited higher responsiveness to methionine restriction were not classified by similar MTAP status or tissue type (Figure S2C).

Of particular interest was the finding that when fold change values of each metabolite were averaged across the 5 cell lines within each group, methionine restriction appeared to have little to no effect on global metabolism (Figure 2E, S2D); in contrast, methionine restriction appeared to have a substantial yet highly variable effect when fold changes were analyzed in each individual cell line (Figure S2E). To further illustrate this finding, we examined alterations in key metabolites of the transsulfuration pathway, which is downstream of the methionine cycle and provides glutathione for intracellular redox reactions (Figure 2F). Independent of MTAP status, some cell lines demonstrated significant alterations throughout this pathway, while others appeared to show no changes as a result of methionine restriction. Furthermore, the alterations that were induced were variable between cell lines; for instance, glutathione was found to increase 2-fold in the MTAP^+/+^ melanoma line UACC-257 upon methionine restriction, while it decreased by almost half in the MTAP^-/-^ bladder line RT4 and was non-significantly altered in the majority of the other cell lines (Figure 2F). Altogether, these findings illustrate that responsiveness to alterations in methionine availability is not predicted by MTAP status, and that accumulation of MTA, which is the key feature of the MTAP metabolic signature, is abrogated by methionine restriction.

### Responsiveness to alterations in other one-carbon nutrient availability is largely MTAP status-independent

Although MTAP status did not appear to be predictive of responsiveness to methionine availability, we further hypothesized that the MTAP-deleted lines may have adapted to the inability to salvage methionine by differentially regulating the utilization of nutrients that feed into and out of the methionine cycle. To test this, we cultured the cell lines in either serine-restricted (~16 uM compared to ~280 uM) media for 24 hours or in cysteine-restricted (~6 uM compared to ~200 uM) media for 6 hours and validated the reduction in levels of the respective amino acid (Figure 3A, S3A). Of note, a shorter culture time for the cysteine restriction experiments was used due to the variable toxicity of 24-hour cysteine restriction across the panel of cell lines (Figure S3B). As observed with methionine restriction, cell lines did not appear to globally respond to either serine or cysteine restriction in an MTAP status-dependent manner and the new metabolic state that arose from altered nutrient availability was independent of MTAP status (Figure 3B).

**Figure 3:**
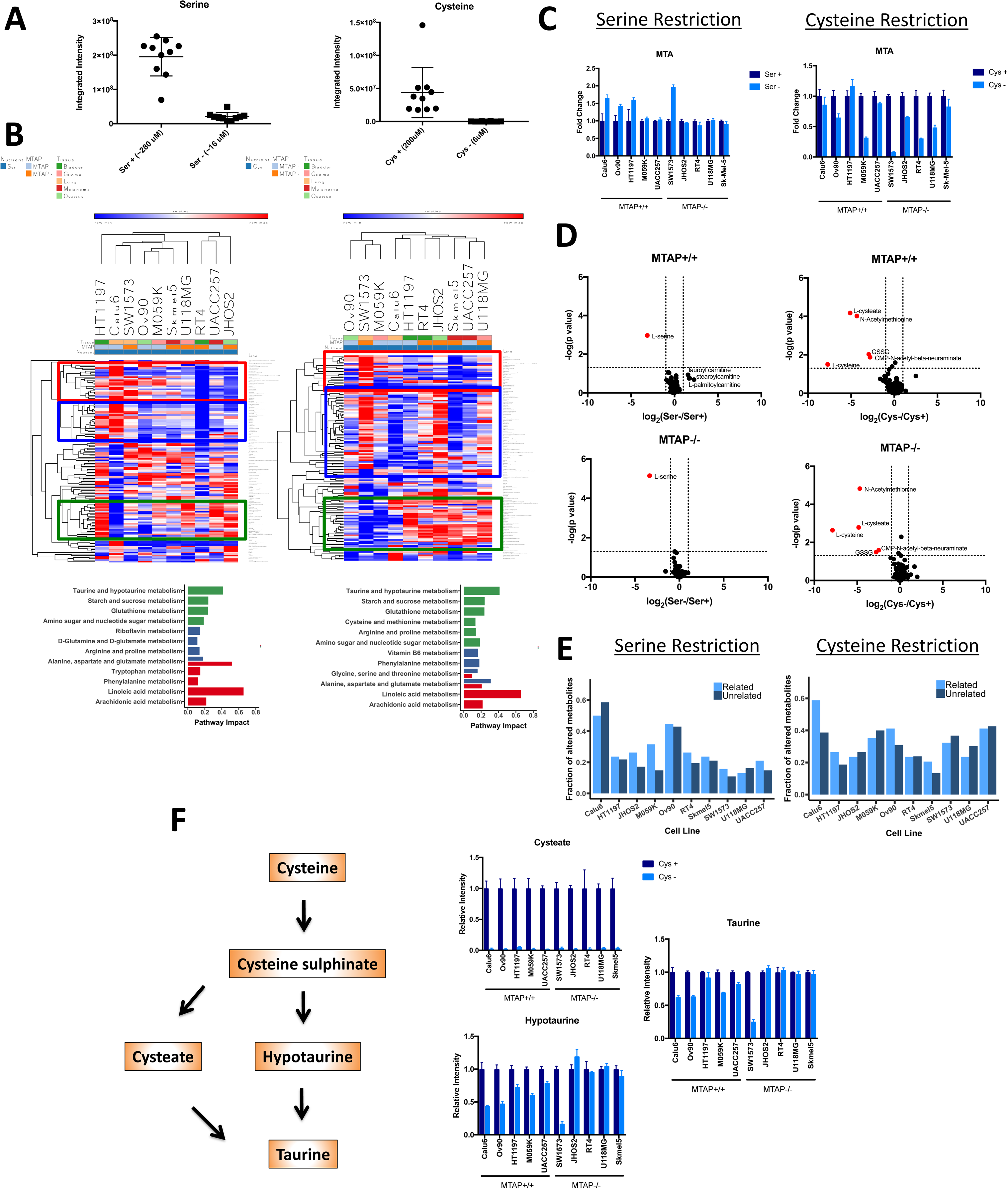
Responsiveness to alterations in other one-carbon nutrient availability is largely MTAP status-independent. (A) Validation of serine and cysteine restriction. (B) Heat map of fold changes (FC) in global metabolite levels upon serine (right) and cysteine (left) restriction, hierarchically clustered by cell lines and metabolites. Top impacted pathways are indicated. (C) FC values of MTA metabolite levels either in complete (280uM Ser, or Ser+, and 200uM Cys, or Cys+) or restricted (~16uM Ser, or Ser-, and 6uM Cys, or Cys-) (D) Volcano plot of FC (Ser-/Ser+, or Cys-/Cys+) values of metabolites, averaged across either MTAP+/+ or MTAP-/- groups. P values obtained using Student’s t-test. (E) Fraction of significantly altered (p<0.05, Student’s t-test) metabolites that are related (i.e. within 5 reactions) or unrelated to serine (left) and cysteine (right). (F) Alterations in relative metabolite levels upon cysteine restriction in metabolites involved in taurine biosynthesis.

Interestingly, serine restriction appeared to induce an increase in MTA levels in three out of the five MTAP^+/+^ lines and in one MTAP^-/-^ line, while producing no significant change in MTA levels in the other lines; conversely, cysteine restriction mostly induced a decrease in MTA levels in both groups (Figure 3C). Similar to what was found with methionine restriction, when the fold changes of metabolites were averaged across the five cell lines for each group, serine restriction was found to have no significant reproducible and general impact on global metabolism (Figure 3D, S3C) although each individual cell line demonstrated responsiveness to the restriction when examined individually (Figure 3E). In contrast, averaging fold changes across the respective cell lines showed both groups responded similarly to cysteine restriction, as evidenced by significant reductions in cysteine-related metabolites such as cysteate and oxidized glutathione (GSSG) (Figure 3D, S3C); however, each cell line still exhibited a unique metabolic response to the restriction as was found in the other nutrient restriction conditions (Figure 3E). Furthermore, tissue of origin was found to account for more variance in the data than MTAP status, although both were found to account for substantially less of the variance compared to heterogeneity between cell lines (Figure S3D,E).

Interestingly, cysteate (an intermediate in taurine biosynthesis) levels were found to be consistently depleted across all cell lines in response to cysteine restriction; closer examination of this pathway demonstrated that with the exception of one cell line (SW1573), taurine and hypotaurine levels tended to be altered less in MTAP^-/-^ cell lines compared to MTAP^+/+^ cell lines (Figure 3F). These findings could imply that MTAP^-/-^ cells adapt to deficiencies in recycling methionine by becoming less dependent on cysteine availability, although MTAP-deleted lines on average did not show decreased cytotoxicity in cysteine-restricted conditions compared to MTAP-positive lines (Figure S3A).

Altogether, these results show in terms of the factors mediating responsiveness to one-carbon nutrient availability, it is largely MTAP-independent and is better predicted by other contextual factors.

### Restoration of MTAP expression produces heterogeneous effects on metabolism

We next sought to determine whether re-expression of MTAP protein in MTAP-deleted cell lines would induce similar effects on both responsiveness to nutrient restriction and on overall metabolism. We ectopically expressed MTAP or a GFP control in four MTAP-deleted cell lines (Figure 4A), and demonstrated that the resulting protein was functional as evidenced by a reduction in MTA levels in MTAP-expressing cells compared to control cells (Figure 4B). Expression of MTAP in these cell lines resulted in changes to both global and methionine metabolism in each of the cell lines; however, only one metabolite (MTA) was significantly altered in all four lines (Figure 4C).

**Figure 4:**
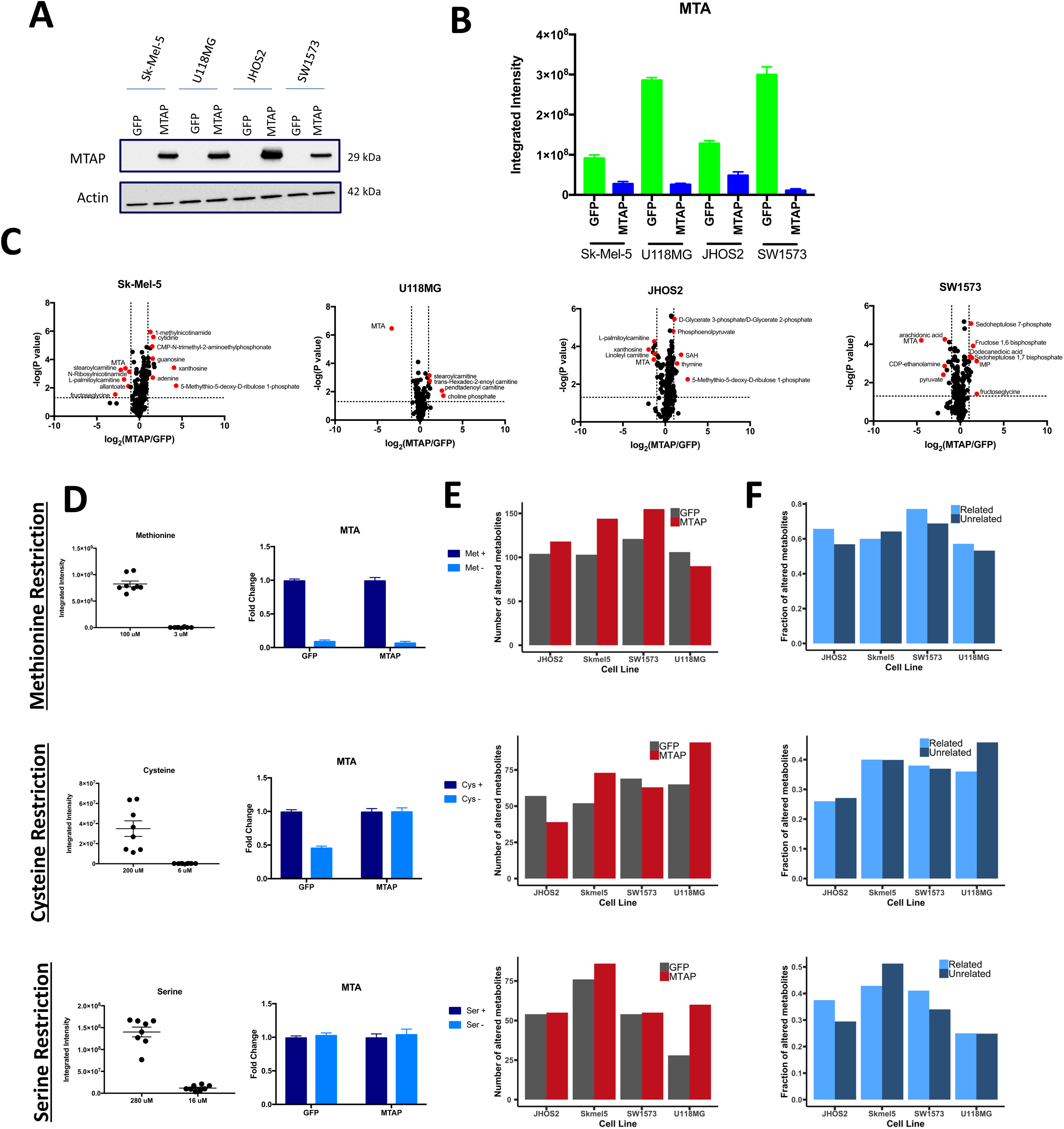
Restoration of MTAP expression produces heterogeneous effects on metabolism. (A) Western blot validation of lentiviral ectopic expression of MTAP (compared to GFP control) in MTAP-/- cell lines. (B) Integrated intensity values of relative MTA metabolite levels in MTAP-or GFP-infected cell lines. (C) Volcano plots of each cell line, showing FC values of metabolites between MTAP-infected and GFP-infected isogenic pairs. P values obtained using Student’s t-test. (D) Validation of nutrient restriction using integrated intensity values of the respective nutrient (left) and FC values of MTA metabolite levels in complete/restricted culture conditions (right). (E) Number of significantly altered (p<0.05, Student’s t-test) metabolites in MTAP-expressing lines compared to GFP controls. (F) Fraction of significantly altered (p<0.05, Student’s t-test) metabolites that are related (i.e. within 5 reactions) or unrelated to the indicated nutrient.

Ectopic MTAP expression did not significantly impact responsiveness of MTA levels to methionine or serine restriction, although cysteine restriction resulted in a significant drop in MTA levels exclusively in MTAP^-/-^ lines (Figure 4D). Interestingly, MTAP re-expression also largely did not appear to significantly alter overall responsiveness to methionine, cysteine, or serine availability compared to controls as evidenced both by the number of metabolites related to methionine (Figure S4A) and to the overall changes in metabolism (Figure 4E, S4B). Additionally, MTAP re-expression also did not seem to increase the proportion of significantly altered metabolites that were related versus unrelated to the respective nutrient that was restricted (Figure 4F). These results imply that while ectopic MTAP expression in *MTAP*-deleted cancer cell lines can induce significant metabolic alterations, these effects are not sufficient to override the responsiveness to nutrient availability that have been adapted by individual cell lines.

### Defining the quantitative impact of MTAP deletion and environmental factors on metabolism

These results so far demonstrate that heterogeneity across cell types and nutrient availability exert dominant effects on metabolism relative to MTAP status. For example, each cell line exhibited marked differences in the number of altered metabolites independent of MTAP status or tissue of origin (Figure 5A). Furthermore, when the number of unique significantly altered metabolites were averaged within MTAP^+/+^ and MTAP^-/-^ groups, we found that the two groups displayed similar degrees of responsivity to methionine restriction while the MTAP^+/+^ cell lines were on average more responsive to serine and cysteine restriction (Figure 5B), as was further confirmed using 3 independent statistical analyses (Student’s t, Kruskal-Wallis, Mann-Whitney U tests) (Figure 5C). An additional comparison of the number of altered metabolites between MTAP^+/+^ and MTAP^-/-^ cell lines demonstrated that each nutrient restriction had a substantially greater impact on global metabolism than *MTAP* deletion by itself (Figure 5D).

**Figure 5:**
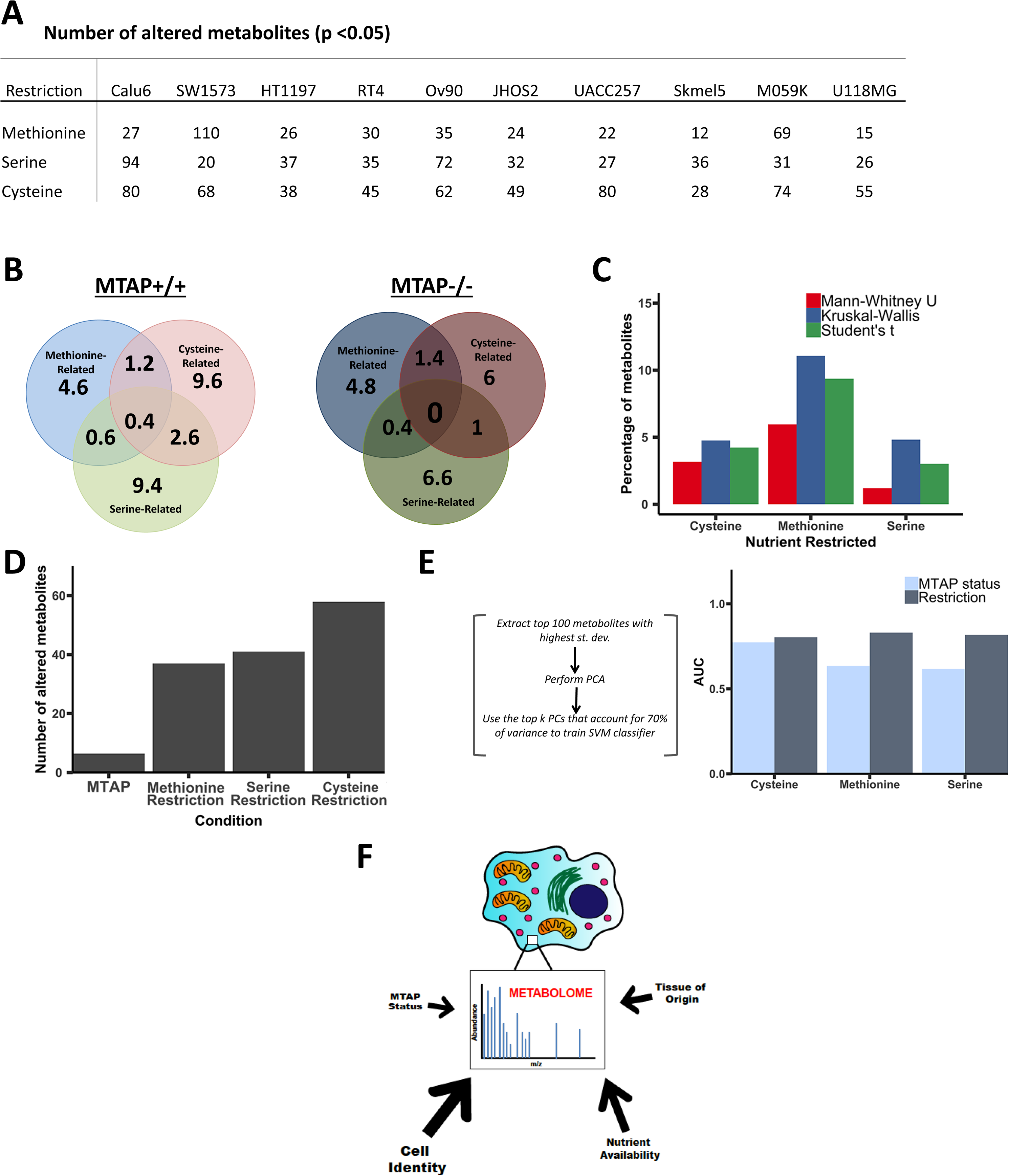
Integration of responsiveness to nutrient restriction determines quantitative impact of MTAP deletion on global metabolic networks. Number of significantly altered (p<0.05, Student’s t-test) metabolites for each cell line under each of the three nutrient restriction conditions. B) Venn diagrams depicting overlap of average number of significantly altered related metabolites within a cell line for MTAP+/+ and MTAP-/- groups. (C) Number of significantly altered metabolites for each nutrient restriction, as determined by three separate statistical analyses as indicated. (D) Number of total unique significantly altered metabolites for each condition indicated. (E) Area under the curve (AUC) of top principal components (PCs) that account for 70% of variance between either MTAP+/+ and MTAP-/- or control and restricted nutrient conditions. (F) Graphical summary illustrating that environment and cell identity shape the metabolome to a greater extent than MTAP status or tissue of origin.

To quantify the relative contribution of MTAP and the availability of each nutrient, we developed a machine learning classifier with a support vector machine using the top 100 metabolites exhibiting the highest standard deviation within each restriction data set (Figure 5E, Methods). First, a PCA analysis was used to generate a vector that accounts for 70% of the variation across all of the samples. This classifier was tested for its ability to discriminate (for both sensitivity and specificity) MTAP status, or cysteine, methionine, or serine availability. In each case of differential nutrient availability, the nutrient status outperformed MTAP status in accounting for the variation across the data. Thus, these collective results demonstrate that while MTAP deletion does confer a reproducible metabolic signature within a single environment, the availability of nutrients related to MTAP (i.e. methionine, serine, and cysteine) has a larger effect on both MTAP-related methionine metabolism and on the global metabolic network, which is further confounded by metabolic variation across cell lines that span different tissues of origin (Figure 5F).

## Discussion

It has recently been proposed that MTAP deletion creates a state of disordered methionine metabolism that creates targetable liabilities, in both metabolism and the epigenetic regulation that is connected to methionine metabolism (8–10, 12, 25); however, the data supporting this model have predominantly come from experiments in cells cultured in complete media. Our results demonstrate that while metabolic signatures of *MTAP* status across diverse cell types are reproducible in such conditions, changes to cellular nutrient availability related to MTAP metabolism reprogram metabolism to greater extents. For instance, we found that the accumulation of MTA, which was essential for mediating the dependency on MTAP, was eliminated when cells were subjected to methionine restriction. Furthermore, metabolic variation across cell types was far greater than the differences observed in a pan tissue analysis of MTAP wild-type and homozygous deleted cells. Other examples of collateral lethality that depend on genetically defined altered metabolism in general, such as the case of malic enzyme or enolase (26, 27), would likely also encounter similar considerations.

There has been a general underlying assumption in the field of cancer metabolism that cancerous cells exhibit similar metabolic phenotypes based on their genetic status (28, 29). Using *MTAP* status and the pathway its enzyme resides in (i.e. methionine metabolism) as a model to quantitatively and systematically study the relevant variables that shape metabolic outcomes, we found that while perhaps expectedly global metabolism was not overall predictive of *MTAP* status, unexpectedly methionine metabolism as well was also shaped more by the nutrients that are used in the one-carbon network and by cell-to-cell variability.

Recently a number of studies have shown that environmental factors influence the outcome of a metabolic pathway (19, 30, 31). For example, a genetic event that generates a tumor, when initiated in different tissues, results in different metabolic programs (19, 20). Our study further develops this concept by showing for the case of MTAP status and methionine, environmental factors can play a dominant role in defining the state of metabolic networks. Altogether, these studies clearly indicate greater complexities in the factors that shape metabolism.

## Author Contributions

Conceptualization, S.M.S and J.W.L.; Cell Culture Experiments, S.M.S.; Metabolomics and Data Analysis, S.M.S. and P.M.; Mathematical Modeling, P.M. and Z.D. All other experiments, S.M.S.; Writing, S.M.S. and J.W.L with essential input from all authors.

## Acknowledgments

This work was supported by the National Institutes of Health (R01CA193256 to J.W.L. and F31CA224973 to S.M.S), as well as by a Graduate Fellowship from the Duke University School of Medicine to S.M.S. We thank the members of the Locasale laboratory for their comments and support, particularly Xiaojing Liu, Michael Reid, and Maria Liberti for their technical assistance.

## Methods

### Cell Culture

Cells were cultured at 37°C, with 5% atmospheric CO_2_ in RPMI-1640 (GIBCO), 10% heat-inactivated fetal bovine serum (FBS), 100 U/mL penicillin, and 100 mg/ml streptomycin. All cell lines were obtained from the American Tissue Culture Collection (ATCC) except UACC-257 (obtained from the National Cancer Institute, National Institutes of Health) and JHOS-2 (obtained from Dr. Andrew Berchuck, Duke University).

### Nutrient Restriction

For all nutrient restriction experiments, cells were plated at a density of 3x105 cells/well in triplicate in a 6-well plate and were allowed to adhere for 24 hours. Conditional media was prepared using RPMI-1640 lacking amino acids, glucose, and glutamine. Dropout media (i.e. Met-, Ser-, or Cys-) was supplemented with 5 mM glucose, 2 mM glutamine, 10% heat-inactivated FBS, 100 U/mL penicillin, 100 mg/mL streptomycin, and 19 out of 20 individual amino acids (excluding either methionine, serine, or cystine) at concentrations found in full RPMI-1640 media. For full media conditions in these experiments, the respective nutrient of interest was individually added back to the media (i.e. Met+, Ser+, or Cys+). For all experiments, 3 technical replicates per culture condition for each cell line were used.

### Cell Viability Assays

For all cell viability measurements, cells were plated at a density of 5x103 cells/well in triplicate in a 96-well plate and were allowed to adhere in full RPMI-1640 media for 24 hours. Conditional media, containing 4 (serine) or 5 (cystine or methionine) increasing concentrations of the respective nutrient, was prepared as previously described. After 24 hours, cells were briefly washed with PBS and then incubated in the conditional media for 24 hours. After 24 hours, the media was aspirated and replaced with 100 μL phenol-red free RPMI-1640 (GIBCO) and 12 mM 3-[4,5-Dimethylthiazol-2-yl]-2,5-diphenyltetrazolium (MTT) (Thermo Fisher Scientific, #M6494). After 4 hours, the MTT-containing media was aspirated and 50 μL DMSO was added to dissolve the formazan. After 5 minutes, absorbance was read at 540nm.

### Lentiviral Transfection and Transduction

HEK-293T cells were plated at a density of 1x106 cells per 10 cm plate in RPMI-1640 (GIBCO) supplemented with 10% heat-inactivated FBS, 100 U/mL penicillin, and 100 mg/mL streptomycin and allow to adhere and reach 70% confluency. 15 *μ*g MTAP (Genecopoeia, EX-A3221-Lv105) or GFP control (Genecopoeia, EX-EGFP-Lv105) plasmid, 10 *μ*g PsPAX2 packaging vector (Addgene, #12260) and 5 *μ*g PMD2.G envelope expressing plasmid (Addgene, #12259) were diluted into 500 *μ*L jetPRIME buffer (Polyplus Transfection, #114-07) and vortexed. Next, 60 *μ*L jetPRIME transfection reagent (Polyplus Transfection, #114-07) was added to the mixture, vortexed for 10 seconds, and left to incubate for 10 minutes at room temperature. The media in the plate was replaced with fresh medium, and the transfection mix was then added to the 10 cm plate dropwise. After 24 hours, the transfection medium was replaced with fresh medium. After an additional 24 hours, the medium was collected and filtered through a 0.45 um filter for virus collection. SW1573, U118MG, JHOS2, and Sk-Mel-5 cells were plated in 10 cm plates, and when they reached 30-50% confluency, -containing media (1:1 with fresh RPMI-1640 media) was added to the plates, along with 4 *μ*g/*μ*L polybrene. After 24 hours, the virus-containing media was removed and replaced with fresh RPMI-1640 media. Cells were incubated with 1 *μ*g/mL puromycin for 48 hours and MTAP expression in the 4 isogenic cell pairs (8 lines total) were verified by immunoblotting.

### Immunoblotting

Samples were homogenized in 100 *μ*L 1X RIPA buffer (VWR International) supplemented with 100 *μ*M phenylmethylsulfonyl fluoride (PMSF), 2 *μ*g/*μ*L aprotinin, 1X phosphatase inhibitor cocktail, and 2 mM dithiotheritol (DTT). Cell lysates were centrifuged at 14,000 rpm for 30 min at 4°C. The resulting supernatant was transferred to a clean tube and a bicinchoninic acid (BCA) assay (Thermo Scientific) was carried out to quantify protein concentration. Protein samples were loaded onto TGX stain-free precast gels (Bio-Rad) and transferred to polyvinylidene difluoride (PVDF) membranes. Membranes were blocked in 5% dry non-fat milk in TBST and incubated in anti-MTAP (Cell Signaling, #4158S) 1:7500 in 5% BSA in TBST and anti-Actin (Thermo Scientific, #MA5-15739) 1:2000 in 5% dry non-fat milk in TBST overnight at 4°C. Membranes were washed 3 times for 10 minutes each with TBST. Horseradish peroxidase-conjugated anti-mouse (Rockland, #610-1302) and anti-rabbit (Rockland, #611-1302), both 1:2000, were used as secondary antibodies in 5% dry non-fat milk in TBST at room temperature for 20 minutes. Membranes were washed 3 times in TBST once more, and then chemiluminescent signals were detected with Clarity Western ECL Detection Kit (Bio-Rad, #1705061). The membranes were then imaged using the ChemiDoc Touch Imaging System (Bio-Rad).

### Metabolite Extraction

Media was quickly aspirated and 1 mL of extraction solvent (80% methanol/water, cooled to -80°C) was added to each well of the 6-well plates, and were then transferred to -80°C for 15 minutes. Plates were removed and cells scraped into the extraction solvent on dry ice. All metabolite extracts were centrifuged at 20,000g at 4°C for 10 min. Finally, the solvent in each sample was evaporated in a speed vacuum, and the resulting pellets were stored in -80°C until resuspension. For polar metabolite analysis, the cell extract was dissolved in 15 *μ*L water and 15 *μ*L methanol/acetonitrile (1:1, v/v) (LC-MS optima grade, Thermo Scientific). Samples were centrifuged at 20,000g for 2 minutes at 4°C, and the supernatants were transferred to Liquid Chromatography (LC) vials.

### Liquid Chromatography

Ultimate 3000 HPLC (Dionex) with an Xbridge amide column (100 x 2.1 mm i.d., 3.5 μm; Waters) is coupled to Q Exactive-Mass spectrometer (QE-MS, Thermo Scientific) for metabolite separation and detection at room temperature. The mobile phase A reagent is composed of 20 mM ammonium acetate and 15 mM ammonium hydroxide in 3% acetonitrile in HPLC-grade water (pH 9.0), while the mobile phase B reagent is acetonitrile. All solvents are LC-MS grade, purchased from Fischer Scientific. The flow rate used was 0.15 mL/min from 0-10 minutes and 15-20 minutes, and 0.3 mL/min from 10.5-14.5 minutes. The linear gradient was as follows: 0 minutes 85% B; 1.5 minutes 85% B, 5.5 minutes 35% B; 10 minutes 35% B, 10.5 minutes 25% B, 14.5 minutes 35% B, 15 minutes 85% B, and 20 minutes 85% B.

### Mass Spectrometry

The Q Exactive MS (Thermo Scientific) is outfitted with a heated electrospray ionization probe (HESI) with the following parameters: evaporation temperature, 120°C; sheath gas, 30; auxiliary gas, 10; sweep gas, 3; spray voltage, 3.6 kV for positive mode and 2.5 kV for negative mode. Capillary temperature was set at 320°C and S-lens was 55. A full scan range was set at 60 to 900 (m/z), with the resolution set to 70,000. The maximum injection time (max IT) was 200 ms. Automated gain control (AGC) was targeted at 3,000,000 ions.

### Peak Extraction and Metabolomics Data Analysis

Data collected from LC-Q Exactive MS was processed using commercially available software Sieve 2.0 (Thermo Scientific). For targeted metabolite analysis, the method “peak alignment and frame extraction” was applied. An input file (“frame seed”) of theoretical m/z (width set at 10 ppm) and retention time of ~260 known metabolites was used for positive mode analysis, while a separate frame seed file of ~200 metabolites was used for negative mode analysis. To calculate the fold changes between different experimental groups, integrated peak intensities generated from the raw data were used. Hierarchical clustering and heatmaps were generated using Morpheus software (The Broad Institute, https://software.broadinstitute.org/morpheus/). For hierarchical clustering, spearman correlation parameters were implemented for row and column parameters. Pathway enrichment analysis was conducted using MetaboAnalyst 3.0 software (http://www.metaboanalyst.ca/faces/home.xhtml); briefly, HMDB IDs from the metabolites that were significantly enriched (greater than log2(2)fold change with p<0.05) were inputted. The pathway library used was *Homo sapiens* and Fishers’ Exact test was employed for over-representation analysis. Other quantitation and statistics were calculated using Graphpad Prism software.

### Statistical Analysis

For the control data from the three nutrient restriction experiments, raw intensities of metabolites were log-transformed (log(x+1)), corrected for batch effect using the *removeBatchEffect()* function in the LIMMA R package (32), and then z-score normalized. Other metabolomic data and fold changes between conditions were also z-score normalized before the following analysis. PCA was performed using the *PCA()* function from the FactoMineR R package (33). Correlation (*r^2^*) between each PC and each of the three categorical variables (MTAP status, Cell Line, Tissue of Origin) was also calculated by the *PCA()* function from FactoMineR. ANOVA was done on each of the three categorical variables and the first 10 PCs using the *aov()* function in R. Unpaired student’s t-test was used for comparison between two groups unless otherwise stated. A P-value<0.05 was considered as statistically significant.

### Machine Learning

The area under the curve (AUC) of the receiver operating characteristic (ROC) curve was computed using the *perfcurve()* function in MATLAB. Classification was done using the MATLAB Classification Learner App. Briefly, for binary classification of MTAP status or nutrient status based on metabolite abundances, the top 100 metabolites with the highest standard deviation in their raw intensities were kept. Average intensities from three replicates were z-score normalized, and then subjecting to PCA with the *pca()* function in MATLAB. The minimal number of PCs that accounted for at least 70% of the variance were used to train the linear SVM model for the classification. The AUC was calculated using 5-fold cross validation.

### Network Analysis

The genome-scale reconstruction of human metabolism, Recon2 (34), was used as the metabolic network model in defining the nutrient related and unrelated metabolites. Metabolites that were five or fewer reactions away from the nutrient (methionine, serine or cysteine) were categorized as ‘related’; otherwise they were categorized as ‘unrelated’. Only enzyme-catalyzed reactions with the highest confidence score were considered. Cofactors associated with more than 100 reactions were also excluded from the analysis.

## Supplementary Figure Legends

**Figure S1: Cancer cell lines exhibiting homozygous deletion of MTAP show altered patterns of metabolite levels.** Heatmap displaying integrated intensity values of all detected metabolites in panel of 10 cell lines cultured in complete media.

**Figure S2: Responsiveness to methionine availability is not predicted by MTAP status.** (A) Fold change (FC) values of amino acids in complete (Met+) and methionine-restricted (Met-) media. (B) Dot plot displaying FC values of individual metabolites, averaged across MTAP+/+ (x-axis) and MTAP-/- (y-axis) cell lines. (C) Comparison of p-values (Student’s t-test) for each principal component (PC) that correspond to each factor of interest, with the highest variance that’s accounted for by the respective factor displayed in the corresponding bar graph and dot plot. (D) Spearman correlation coefficients calculated for the similarity in responsiveness to methionine restriction between each cell line, for all metabolites or specifically methionine-related metabolites. (E) Volcano plots of FCs for each individual cell line (top line displaying MTAP+/+ lines, bottom line displaying MTAP-/- lines).

**Figure S3: Responsiveness to alterations in other one-carbon nutrient availability is largely MTAP status-independent.** (A) Viability of each cell line cultured for 24 hours in media containing the respective concentration of cysteine. P values obtained using Student’s t-test. (B) Fold change (FC) values of amino acids in complete (Ser+ or Cys+) and nutrient-restricted (Ser-or Cys-) media. (C) Dot plot displaying FC values of individual metabolites, averaged across MTAP+/+ (x-axis) and MTAP-/- (y-axis) cell lines. (D) Variance attributed to the indicated factor, along with the corresponding PC that displays the highest variance between groups within each factor (displayed as both bar graphs and dot plots). (E) Spearman correlation coefficients calculated for the similarity in responsiveness to methionine restriction between each cell line, for all metabolites or specifically serine- or cysteine-related metabolites. (F) Volcano plots of FCs for each individual cell line (top line displaying MTAP+/+ lines, bottom line displaying MTAP-/- lines).

**Figure S4: Restoration of MTAP expression produces heterogeneous effects on metabolism.** (A) Example of heterogeneous responsiveness to methionine restriction between isogenic lines exhibiting restoration of MTAP expression, as indicated by FC in metabolites involved in the transsulfuration pathway. (B) Dot plot of PCs that display highest variance between the indicated groups, either complete vs restricted media conditions (top) or MTAP+/+ vs GFP control (bottom), with each dot corresponding to individual samples.

